# Crystal structure and biochemical activity of the macro domain from rubella virus p150

**DOI:** 10.1101/2023.07.28.550942

**Authors:** Guido A. Stoll, Liao Zhang, Yorgo Modis

**Author notes:** Correspondence and requests for materials should be addressed to Y.M. at MRC-LMB, Francis Crick Avenue, Cambridge CB2 0QH, United Kingdom.

## Abstract

Rubella virus remains a global health threat. Rubella infections during pregnancy can cause serious congenital pathology and no antiviral treatments are available. Rubella virus encodes a nonstructural polyprotein with RNA polymerase, methyltransferase, and papain-like cysteine protease activities, along with a putative macro domain of unknown function. Macro domains bind ADP-ribose adducts, a post-translational modification that plays a key role in host-virus conflicts. Some macro domains can also remove the mono-ADP-ribose adduct or degrade poly-ADP-ribose chains. Here, we report high-resolution crystal structures of the macro domain from rubella virus nonstructural protein p150, with and without ADP-ribose bound. The overall fold is most similar to macroD-type macro domains from various nonviral species. The specific composition and structure of the residues that poised for catalysis or coordinate ADP-ribose in the rubella virus macro domain are most similar to those of macro domains from alphaviruses. Isothermal calorimetry and enzymatic assays show that the rubella virus macro domain binds ADP-ribose in solution and has mono-ADP-ribosylhydrolase (de-MARylation) activity.

**IMPORTANCE:** Our work demonstrates that, like alpha- and coronaviruses, rubiviruses encode a mono-ADP-ribosylhydrolase with a structurally conserved macro domain fold to counteract MARylation by PARPs in the host innate immune response. Our structural data will guide future efforts to develop novel antiviral therapeutics against rubella or infections with related viruses.

## INTRODUCTION

Rubella virus (RuV), from the *Rubivirus* genus (family *Matonaviridae*), causes rubella, also known as German measles. Rubella is usually a mild disease but infection in the first trimester of pregnancy can result in miscarriage or infants with congenital rubella syndrome (CRS) (1). CRS symptoms include cataracts, deafness, and congenital hearth or brain defects. Rubella virus only infects humans, via the respiratory route, but two recently discovered rubiviruses, ruhugu and rustrela viruses, infect multiple other mammalian species (2). A single vaccine dose, usually as part of the measles, mumps, and rubella (MMR) vaccine, is 95% effective to prevent rubella, but no antiviral treatments are available (1). Rubella remains a common infection worldwide, with 100,000 annual CRS cases (1).

Rubella virus is a member of the alphavirus supergroup, referred to since 2019 as the *Alsuviricetes* class (3), which includes rubiviruses, alphaviruses, hepatitis E virus, and nine other families of viruses that mostly infect plants. Rubella virus is a positive-strand enveloped RNA virus that forms pleiomorphic virions 50-90 nm in length (4, 5). Rubella virus has a 10-kilobase RNA genome with exceptionally high GC content (70%) (4, 6). The virus encodes two polyproteins, one structural and one nonstructural (4). The nonstructural protein, p200, is autoproteolytically cleaved early in infection into the products necessary for viral replication, the p150 and p90 proteins (7). p90 contains the helicase and RNA-dependent RNA polymerase (RdRp) activities required for genome replication (8). p150 contains a methyltransferase domain and a papain-like cysteine protease domain (9, 10). The p150 papain-like protease in has a similar overall structure and catalytic core as the papain-like proteases from SARS-CoV-2 and foot-and-mouth disease virus (10). The papain-like protease is the only protease encoded by rubella virus and is responsible for processing of the viral polyprotein (4, 7, 9).

The papain-like proteases from *Alsuviricetes* and coronaviruses are flanked by a macro domain (9, 11). Macro domains bind ADP-ribose, an important post-translational modification, and some of its metabolites or derivatives (12). Macro domains may catalyze the removal of the mono-ADP-ribose (MAR) adduct, usually to a glutamate or aspartate residue, or the degradation of poly-ADP-ribose (PAR) chains (13, 14). Crystal structures of the macro domains from alpha- and coronaviruses, including SARS-CoV-2, identify the ADP-ribose binding pockets of viral macro domains (15–26). The conserved macro domains from alphaviruses, coronaviruses, and hepatitis E virus are mono-ADP-ribosylhydrolases, or de-MARylases (15, 23–27). Some viral macro domains can also hydrolyze PAR chains, albeit inefficiently (15, 27). Most of the conserved residues that coordinate ADP-ribose in viral macro domains are required for de-MARylase activity (15). The macro domain from mouse hepatitis virus (MHV), a coronavirus, is responsible for liver pathology (28). Conversely, MARylation of aspartate, glutamate, and lysine side chains by a set of MARylases from the PARP family, including PARP10 and PARP14, inhibits translation and alphavirus replication (15, 29–31). These PARPs are interferon-stimulated genes (ISGs) (29) and are evolving rapidly under strong recurrent positive selection (32). Addition and removal of MARylation by cellular PARP catalytic domains and viral macro domains, respectively, is therefore an important component of host-virus conflict (15, 30, 32). Viral macro domains and their active sites are proving to be attractive targets for antiviral therapeutics, as illustrated by a recent structure-based study identifying small molecules that target the conserved macro domain (macro 1) from SARS-CoV-2 (33).

The structure and biochemical properties of the putative macro domain from rubella virus remain unknown. Here, we report high-resolution crystal structures of the rubella virus p150 macro domain, with and without ADP-ribose bound. The overall fold is most similar to macroD-type macro domains from various viral and nonviral species. The detailed composition and structure of the residues poised for catalysis or coordinating ADP-ribose in the rubella virus macro domain are most similar to those of macro domains from alphaviruses. We show that the rubella virus macro domain binds ADP-ribose in solution and has de-MARylation activity. Our work demonstrates that, like alpha- and coronaviruses, rubiviruses encode a de-MARylase with a structurally conserved macro domain fold to counteract MARylation by PARPs in the host innate immune response. Our structural data will guide future efforts to develop novel antiviral therapeutics against rubella or infections with related viruses.

## RESULTS

### Crystal structure of the RuV p150 macro domain

We first determined the crystal structure of the macro domain from rubella virus p150 (RuV macro), spanning residues 805-983 of p150 (UniProt G3M8F4). The structure was determined at 1.7 Å resolution by automated molecular replacement with MrBUMP (34, 35) using an atomic search model based on the human protein-proximal ADP-ribosyl-hydrolase MacroD2 structure (PDB 4IQY) (36). See **Table 1** for crystallographic data collection, refinement, and validation statistics. The overall fold is typical for a macro domain: a central 7-stranded β-sheet surrounded by five peripheral α-helices (**Fig. 1A**). The most structurally similar proteins identified with DALI (37) were macro domains from nonviral species including bacteria, archaea and vertebrates: *Oceanobacillus iheyensis* macroD (Z-score 23.7, rmsd 2.6 Å, PDB 5LAU and 5L9K (38)); *Archaeoglobus fulgidus* AF1521 macro (Z-score 23.7, rmsd 2.2 Å, PDB 1HJZ (39) and 2BFQ (12)); and human PARP14 macro 1 (Z-score 23.5, rmsd 2.1 Å, PDB 3Q6Z (31)). However, the overall fold of RuV macro also closely resembles those of macro domains from coronaviruses and alphaviruses, for example SARS-CoV-2 macro 1 (Z-score 21.0-21.6, rmsd 2.2-2.3 Å, PDB 6WOJ, 6W02, 6YWL and 7TX5 (20, 21, 23, 24)) and Getah virus macro (Z-score 20.5, rmsd 2.6 Å, PDB 6R0F and 6R0G (22)).

**Fig. 1.**
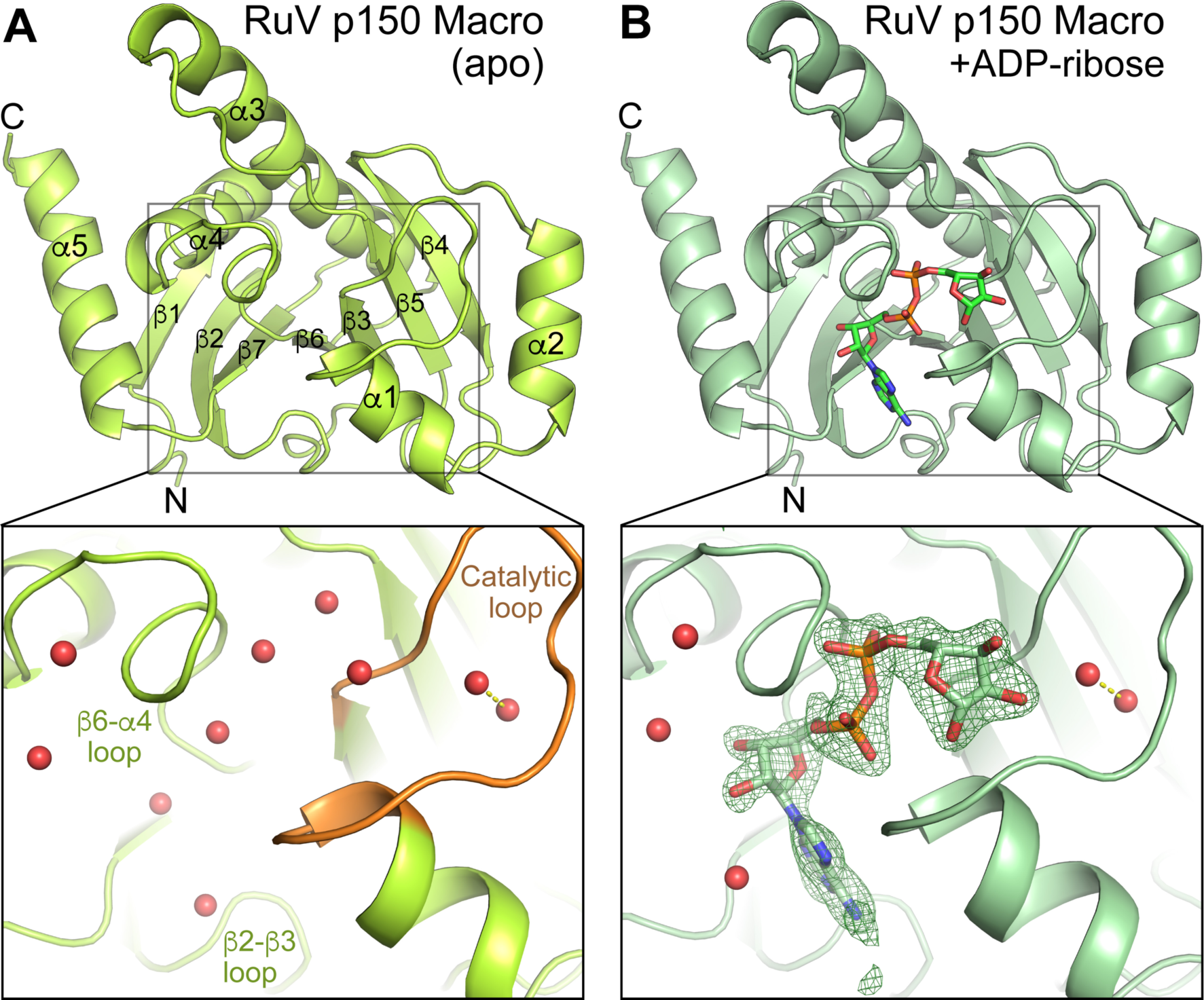
Crystal structures of the macro domain from rubella virus (RuV) p150. **(A)** Structure of RuV macro alone, with the catalytic loop (β3-α1 loop) in orange. Red, water molecules. Dashed line, hydrogen bond. The three substrate-coordinate loops are labeled. **(B)** Structure of RuV macro with ADP-ribose (ADPr) bound. Closeup panels show ADPr binding pocket. The green mesh is a polder map calculated with the ADPr molecule omitted (more specifically, an *F*_obs_ – *F*_calc_ Fourier difference map was calculated with ADPr omitted from the model and the bulk solvent excluded from the omitted region, as implemented in Phenix (43)). An isomesh contour level of 4.2 0 in PyMol (Schrödinger, LLC) was used. See Table 1 for crystallographic data collection, refinement, and validation statistics.

**Table 1.**
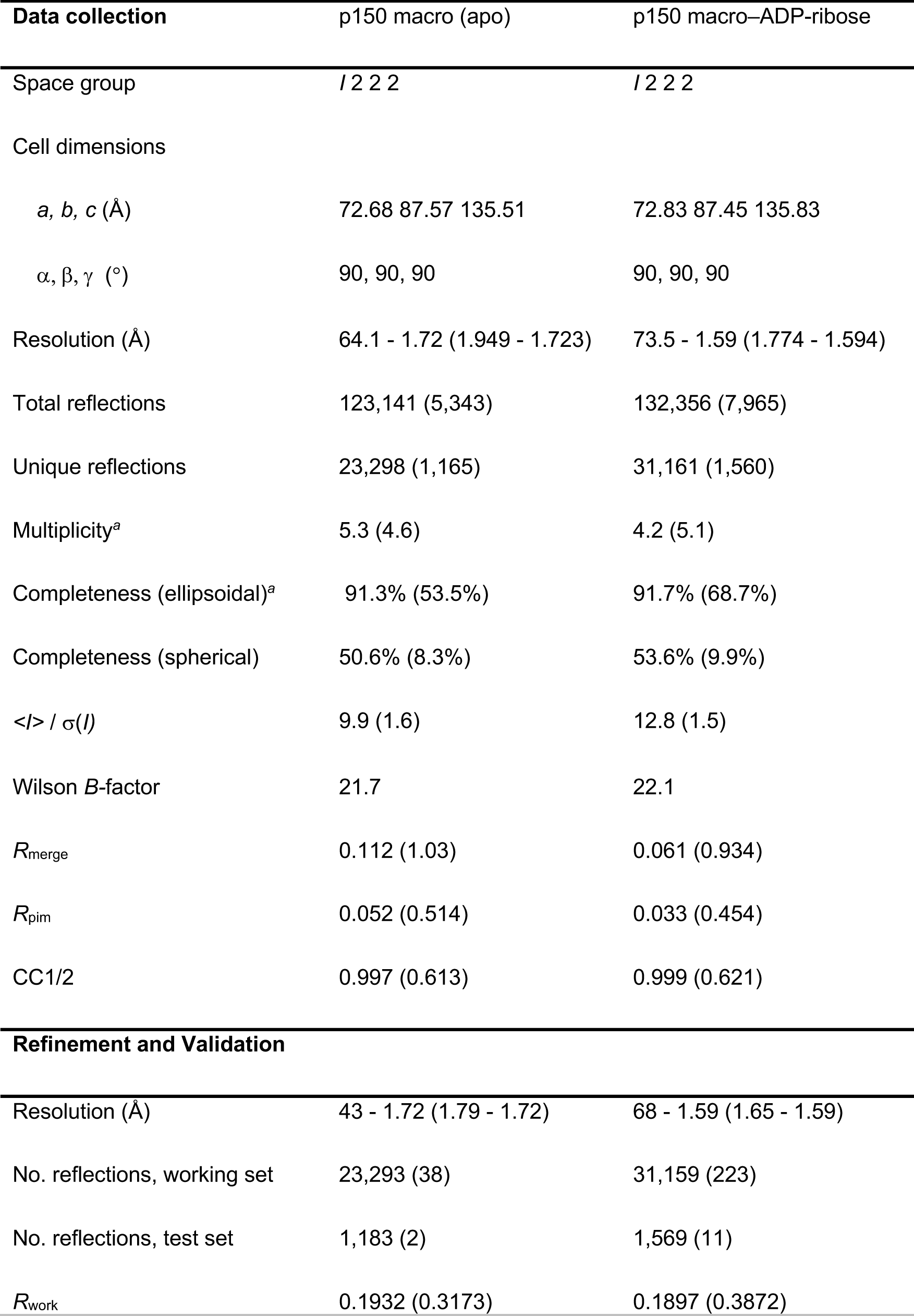
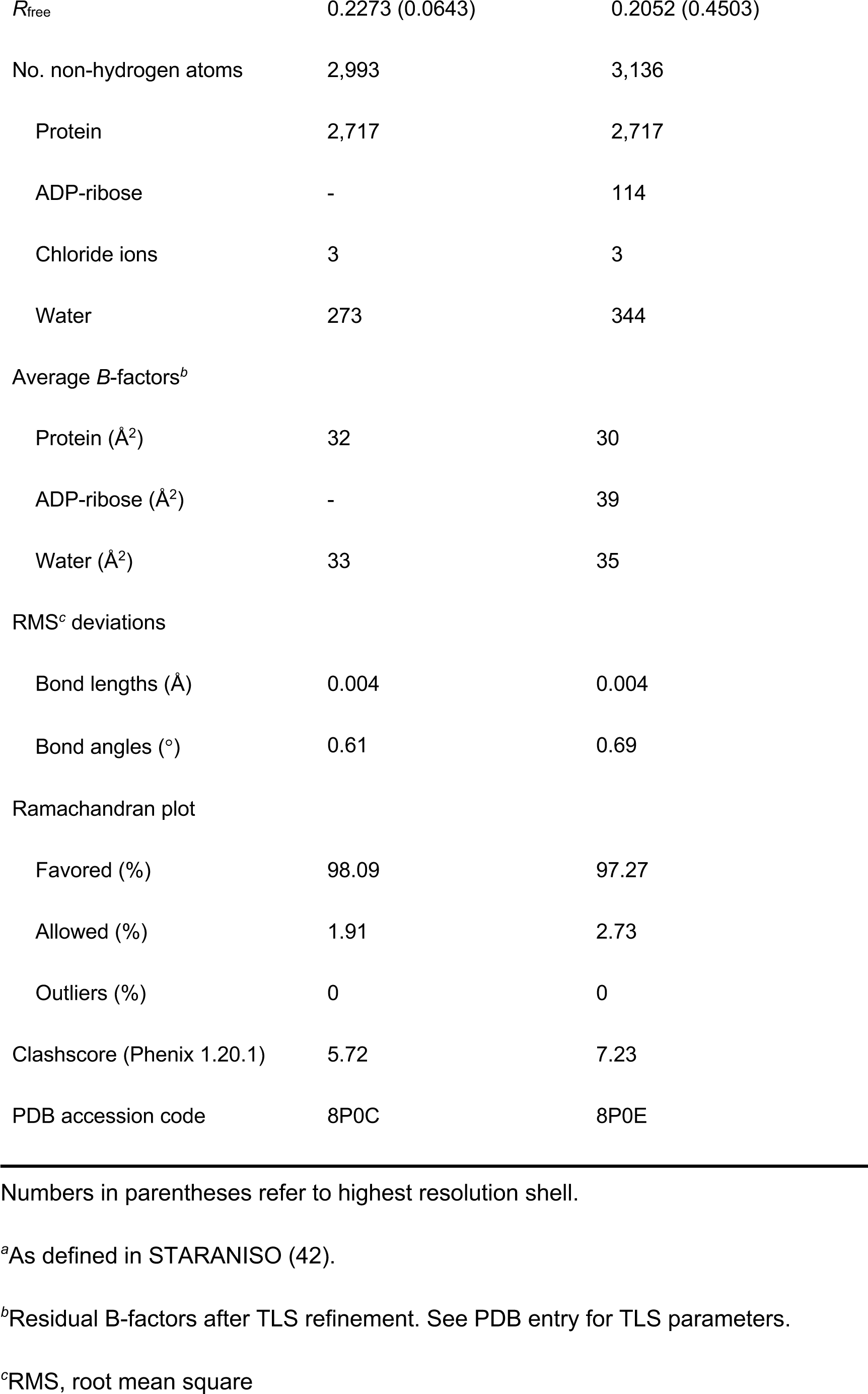
Crystallographic data collection, refinement, and validation statistics.

### Structure of RuV macro with ADP-ribose bound

Since the macro domains with the most similar structures to RuV macro could all be crystallized with ADP-ribose bound, we soaked RuV macro crystals in a solution containing 2 mM ADP-ribose and hence obtained a structure with ADP-ribose bound. The ADP-ribose molecule is in in the β-anomeric configuration and binds to RuV macro in a similar manner to other macro domains (**Fig. 1B**). The binding pocket is formed primarily by the β3-α1 loop, also known as the catalytic loop or Loop 1, the β6-α4 loop, and the β2-β3 loop. The ADP-ribose molecule forms two hydrogen bonds and several hydrophobic contacts with side chains that are broadly conserved in macro domains from different viral, bacterial, and vertebrate species (**Figs. 2 and 3**). One of the conserved hydrogen bonds is between an aspartate in the β2-β3 loop (Asp23 in RuV macro) and the amide group in the adenine moiety of ADP-ribose (**Fig. 2A**). The second hydrogen bond is between the second asparagine in the catalytic loop (Asn39 in RuV macro) and the 3’ or 2’ oxygen atom in the ribosyl (distal ribose) moiety of ADP-ribose. The hydrophobic contacts are with conserved aliphatic side chains in all three of the substrate-coordinating loops. In RuV macro, these aliphatic residues are Ile24 in the β2-β3 loop, Ala37 and Val48 in the catalytic loop, and Pro132 and Val138 in the β6-α4 loop (**Fig. 2A**). Additionally, a conserved aromatic side chain in the β6-α4 loop (Tyr139 in RuV macro) forms a hydrophobic contact with the 4’ and 5’ carbon atoms in the distal ribose. Also conserved are three water molecules directly coordinating the ADP-ribose molecule, forming hydrogen bonds with the α- and β-phosphate, the distal ribose, and multiple residues the catalytic loop (**Fig. 2A-F**). One of these water molecules, positioned between the α-phosphate and distal ribose, has been proposed to mediate substrate-assisted catalysis in human MacroD2 (36) (see below). The conserved residues and water molecules involved in ADP-ribose binding in the RuV macro structure, and the interatomic contacts they form are shown in **Fig. 3**.

**Fig. 2.**
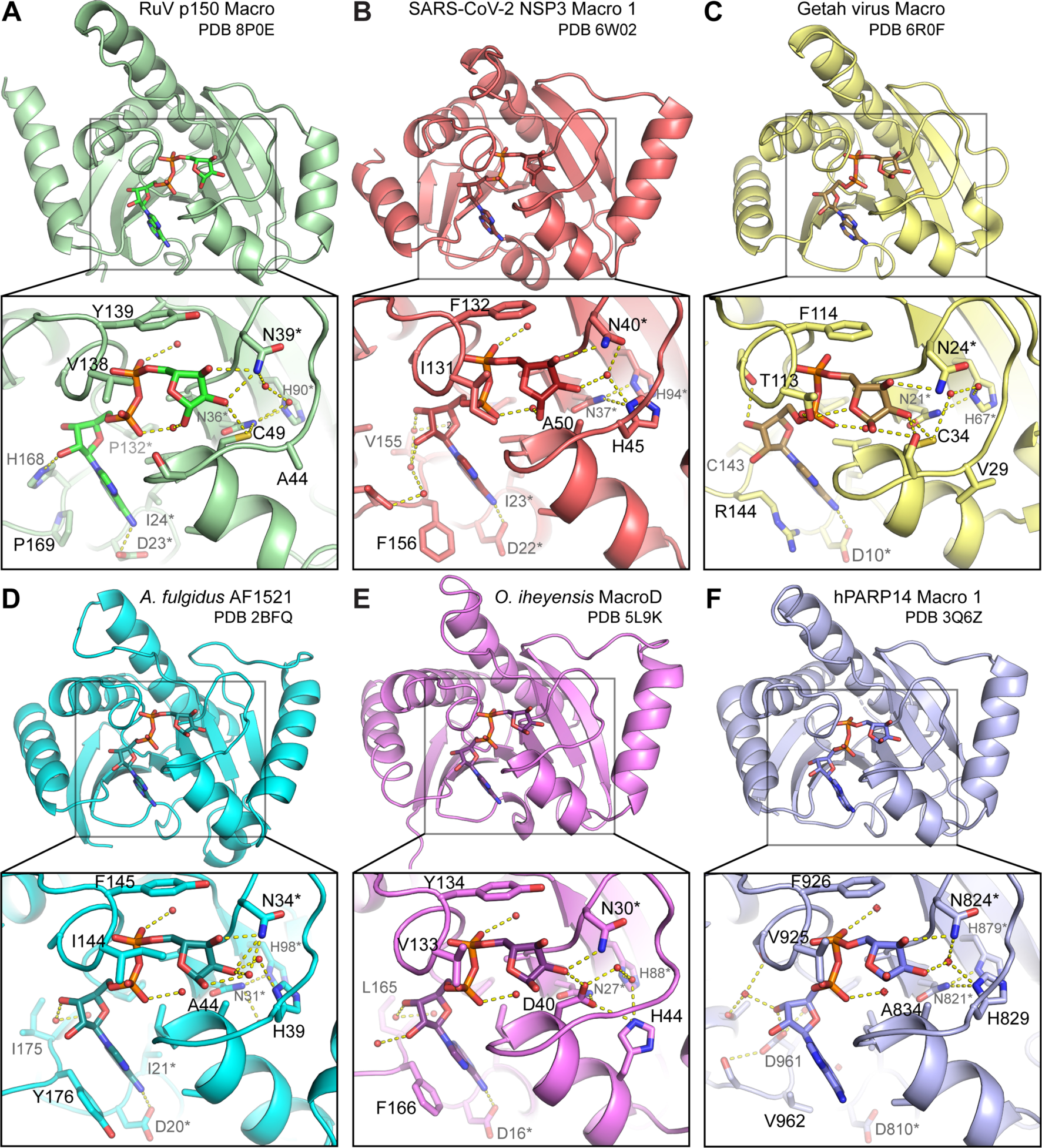
Comparison of the RuV macro structure to macro domains with similar structures. Overviews and closeups of the ADP-ribose (ADPr) binding pockets of: (**A**) RuV macro, (**B**) SARS-CoV-2 NSP3 macro 1, (**C**) Getah virus macro, (**D**) *Archaeoglobus fulgidus* AF1521 macro**, (E**) *Oceanobacillus iheyensis* macroD, (**F**) human PARP14 macro 1. Side chains interacting with ADPr are shown; asterisks denote conserved residues. Dashed lines represent hydrogen bonds.

**Fig. 3.**
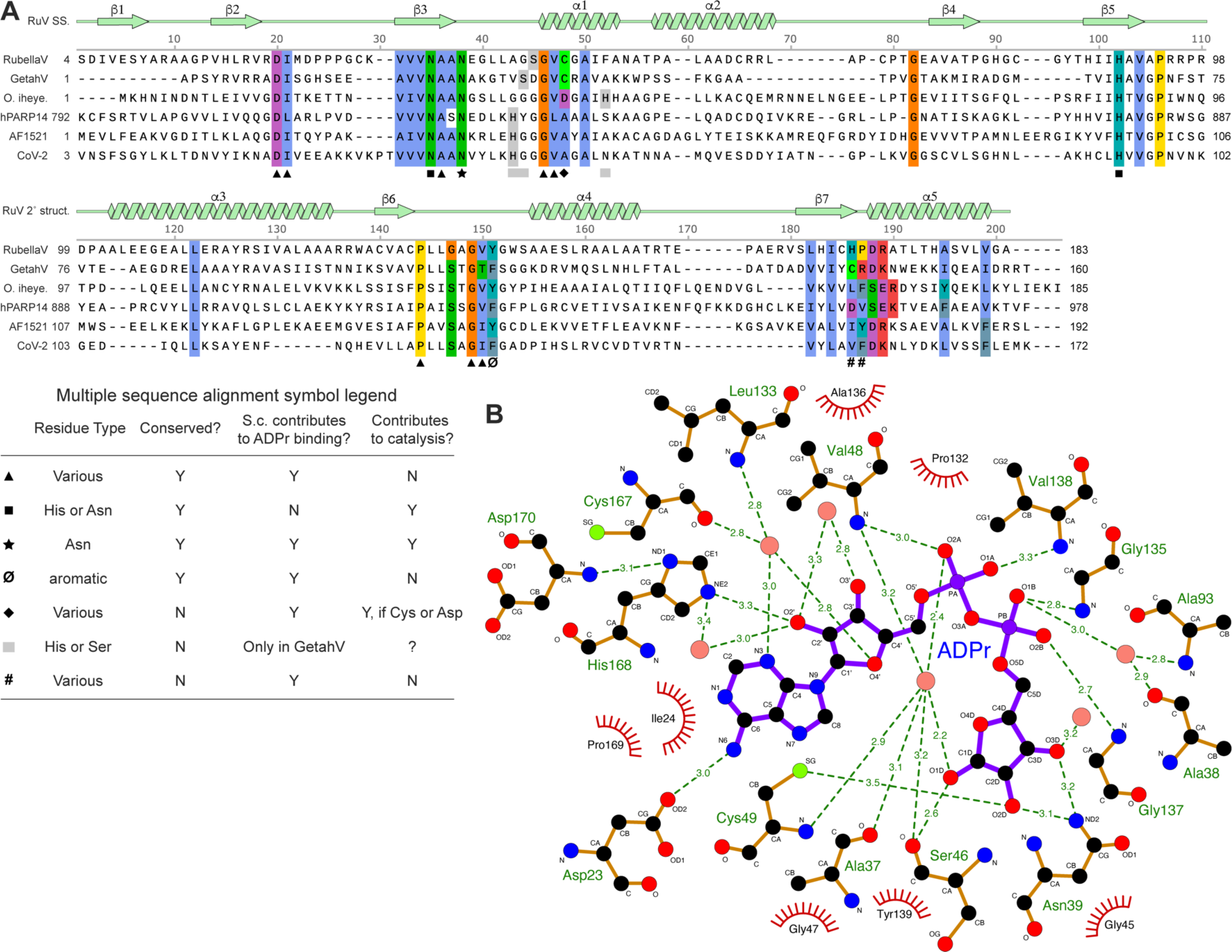
Sequence conservation and interatomic contacts in the RuV macro ADP-ribose (ADPr) binding pocket. (**A**) Structure-based protein sequence alignment of RuV macro and the structurally similar macro domains shown in Fig. 2: Getah virus macro (GetahV), *O. iheyensis* macroD (O. iheye.), human PARP14 macro 1 (hPARP14), *A. fulgidus* AF1521 macro, and SARS-CoV-2 NSP3 macro 1 (CoV-2). The secondary structure of RuV macro is shown above. Residues forming interactions with ADPr and selected conserved residues near the ADPr binding pocket are colored by residue type. Residues forming key contacts with ADPr or potentially contributing to catalysis are denoted with black symbols – see table (lower left) for legend. (**B**) Schematic of the RuV macro residues that form hydrogen bonds (green dashes, with bond length in Å) and hydrophobic contacts (red arcs) with ADPr. Pink, water molecules. S.c., side chain. Based on output from LigPlot+ v.2.2.8 (45).

In addition to the contacts with conserved side chains listed above, multiple hydrogen bonds with main chain atoms of residues in all three substrate-coordinating loops contribute to ADP-ribose binding (**Fig. 3**). Many of these residues (residues 24, 46, 48, and 135-139 in RuV macro) lack sequence conservation, consistent with the ligand contacts involving main chain atoms. Furthermore, unconserved side chain residues in the β7-α5 loop of macro domains from different species (residues 168-169 in RuV macro) form various types of contacts with ADP-ribose, including hydrogen bonds, hydrophobic interactions, and ρε-stacking interactions with the adenosine moiety of ADP-ribose (**Fig. 2A-F**).

### Conserved and novel features in the putative RuV macro active site region

The ADP-ribose binding pocket is broadly conserved in macro domains but there are differences in the composition and structure of the active sites of macro domains from different classes or with different catalytic activities. The closest structural homologs of RuV macro are macroD-type de-MARylases that remove ADP-ribosylation of glutamate or aspartate residues, namely *O. iheyensis* macroD (38), *A. fulgidus* AF1521 macro (39), SARS-CoV-2 macro 1, and Getah virus macro (22) (**Fig. 2A-E**). The defining active site signatures shared by these and other macroD-type domains are Nx(6)GG[V/L/I] and G[V/I/A][Y/F]G motifs in the catalytic and β6-α4 loops, respectively, and water molecules with catalytic potential at conserved positions coordinating the α- and β-phosphate groups (13, 36, 38). The human PARP14 macro 1 domain, though previously classified as macroH2A-like (13), contains all the macroD active site signatures and has comparable structural homology to RuV macro (**Fig. 2F**). The putative active site of RuV macro contains all of the macroD-type signatures including the catalytic asparagine, Asp39 in RuV macro (**Fig. 2A**), except for a serine substitution in the first macroD motif (see below). The RuV macro active site lacks lysine and glutamate residues, which are required for catalysis in macro domains from the ALC1-like (TARG1) and PARG-like classes, respectively (14, 40). Thus, based on the sequence and structure of RuV macro active site residues, we conclude that RuV macro is a member of the macroD-type class.

Although RuV macro contains the macroD-type active site signatures, some atypical substitutions are present in the putative active site. Most notably, RuV macro has a cysteine in the catalytic loop, at position 49 at the N-terminal end of helix α1 (**Figs. 2A and 3A**). An aspartate is most commonly found at this position in the closest macroD homologs, including *O. iheyensis* macroD, in which the aspartate (Asp40) contributes to substrate binding and catalysis (**Figs. 2E and 3A**) (38). However, an aspartate at this position is not absolutely required for de-MARylation activity (36, 38). A cysteine is also present at the equivalent position in the other rubiviruses (ruhugu and rustrella viruses), alphaviruses (25, 26), and in a minority fraction of the most similar protein sequences to RuV macro, mostly from microbiotal species of bacteria. These bacterial species include high GC Gram-positive bacteria such as *Bifidobacterium longum*, which is worth noting given the exceptionally high GC content of the rubella virus genome (4, 6). Crystal structures of the macro domain from Getah virus, from the alphavirus genus, show that the equivalent cysteine (Cys34 in Getah virus) can form a covalent adduct with ADP-ribose (22). In the RuV macro structure, the Cys49 sulfhydryl forms hydrogen bonds with ADP-ribose and a conserved structured water molecule that is also hydrogen bonded to the second asparagine in the catalytic loop, Asn34 in RuV macro (**Fig. 2A**). Together, the substrate interactions of Cys34 in Getah virus macro and Cys49 in RuV macro suggest that a cysteine in this position in the catalytic loop may contribute directly to catalysis (22) in the macro domains of alpha- and rubiviruses.

Another potentially important atypical substitution in the catalytic loop of RuV macro is a serine at residue 46 instead of a glycine in the first macroD signature motif, resulting in the sequence Nx(6)**S_46_**GV instead of the canonical Nx(6)GGV (**Fig. 3A**). Similarly, Getah virus macro has an unusual serine substitution in the preceding position, Ser30. The side chain of Ser30 forms contributes significantly to ADP-ribose binding, forming two hydrogen bonds with the distal ribose (**Fig. 2C**) (22). The Ser46 side chain in RuV macro is similarly oriented, pointing towards the distal ribose of the bound ADP-ribose (**Fig. 2A**), although the minimum distance between the hydroxyls of the serine and the distal ribose, at 4.5 Å, is greater in RuV macro than in Getah virus macro (2.8 to 3 Å). Overall, in comparison to the available ADP-ribose-bound macro domain structures we conclude that the RuV macro structure is most similar in its putative active site composition and organization to the structures of alphavirus macro domains.

### ADP-ribose specifically binds and stabilizes RuV macro in solution

We have shown above that the structure and ADP-ribose binding mode of RuV macro are similar to those of other catalytically active macroD-type domains, but the ADP-ribose binding affinity and catalytic activity of RuV macro in solution remain unknown. We therefore performed solution-based ligand binding assays with RuV macro and ADP-ribose. Isothermal titration calorimetry (ITC) showed that RuV macro binds ADP-ribose with a dissociation constant (*K*_D_) of 58.1 ± 5.4 µM (**Fig. 4A**). The shape of the isotherm curve was indicative of specific, saturable binding. As expected from the structure of RuV macro in complex with ADP-ribose, the protein:ADP-ribose binding stoichiometry was 1:1. Moreover, differential scanning fluorimetry (DSF) using the intrinsic fluorescence of RuV macro as the readout revealed a clear thermal stabilization of the protein on binding ADP-ribose, with an increase in melting temperature of 5.3°C upon saturation of the ADP-ribose binding site (**Fig. 4B**).

**Fig. 4.**
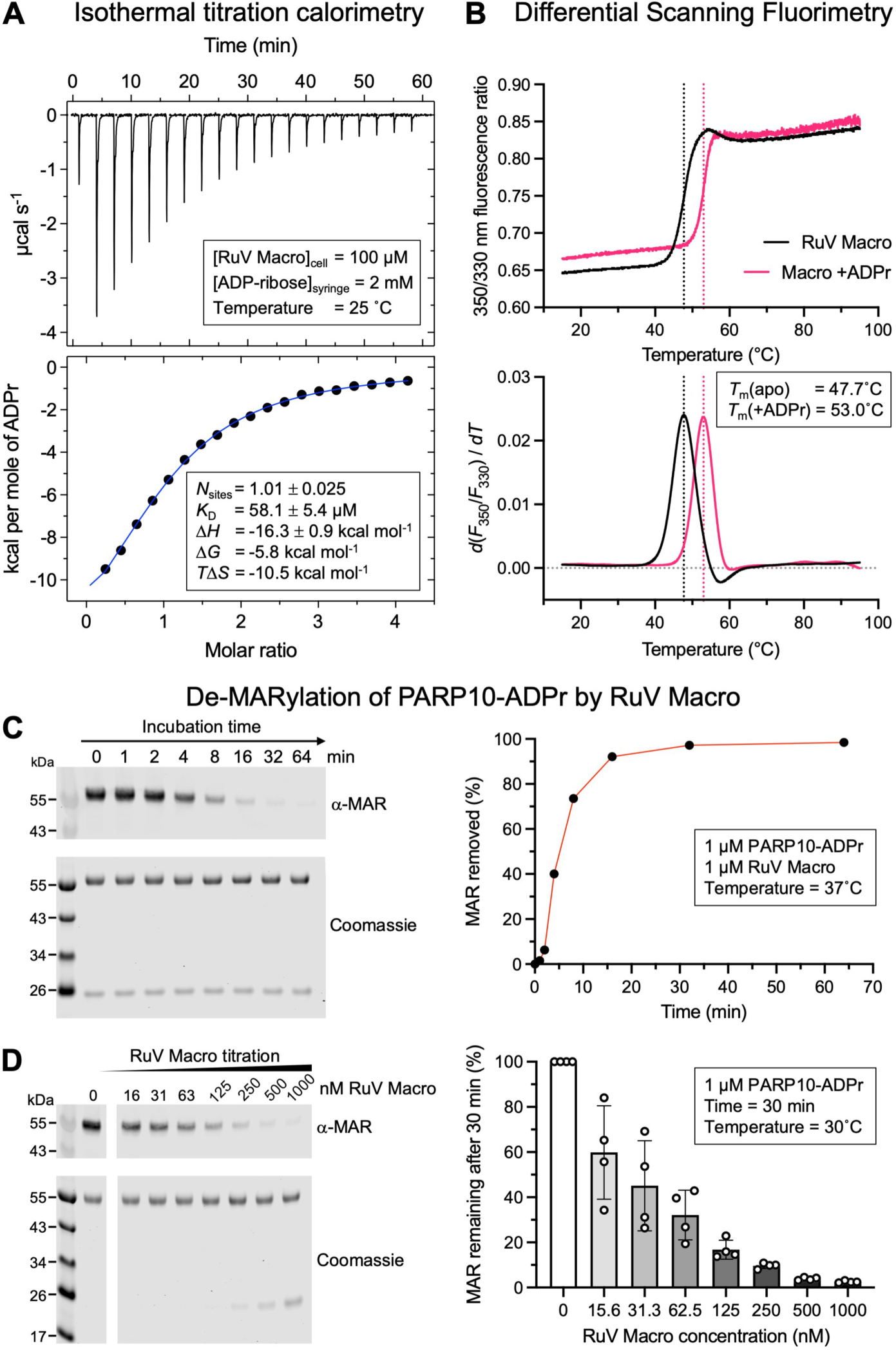
ADP-ribose (ADPr) binding and mono-ADP-ribosylhydrolase assays. (**A**) Isothermal titration calorimetry (ITC) binding assay with RuV macro in the sample cell and ADPr as the injectant. (**B**) Differential scanning fluorimetry (DSF) of RuV macro with and without 5 mM ADPr. Intrinsic protein fluorescence at 330 nm and 350 nm was measured and the fluorescence ratio plotted as a function of temperature. The temperature value at the peak of the first derivative of the fluorescence ratio versus temperature function was taken as the melting temperature (T_m_).

### The RuV macro has mono-ADP-ribosylhydrolase activity

To determine whether RuV macro has mono-ADP-ribosylhydrolase (de-MARylase) activity, like other macroD-type structures, we performed a de-MARylation assay following a previously established protocol (23). Recombinant PARP10 catalytic domain was purified and incubated with NAD+ to allow it to MARylate itself. The ability of RuV macro to de-MARylate the auto-MARylated PARP10 protein was then assayed as a function of time and RuV macro concentration. This de-MARylation assay clearly showed that RuV macro was able to remove mono-ADP-ribosyl adducts from PARP10 (**Fig. 4C,D**). The extent, rate, and enzyme concentration dependence of PARP10 de-MARylation by RuV macro were similar to those reported for SARS-CoV-2 macro 1 (23). Hence, RuV macro is a catalytically active macroD-type macro domain with de-MARylase activity.

## DISCUSSION

We have shown here that the predicted macro domain from rubella virus p150 (RuV macro) has a similar structure and active site to macroD-type mono-ADP-ribosylhydrolases (de-MARylases). The RuV macro active site contains all of the macroD-type signatures except for a serine substitution in the first macroD motif. A detailed comparison with the available macro domain structures shows that the active site composition and organization of RuV macro are most similar to those of macro domains from alphaviruses, and to a lesser extent coronaviruses. RuV macro binds ADP-ribose in a similar manner to other macro domains. Solution-based ligand binding assays show that ADP-ribose binds to a single specific site on RuV macro, resulting in a significant thermal stabilization of the fold. Moreover, we have shown that RuV macro has de-MARylase activity.

The RuV macro active site contains all the hallmarks of a catalytically active de-MARylase and a few atypical features. The second asparagine in macroD motif, Asn39 in RuV macro, is arguably the most important residue for catalysis based on its position in the structure and degree of conservation. Cys49, in the catalytic loop and coordinating ADP-ribose, is also likely to play a role in catalysis. Replacing the more common aspartate, a cysteine is present in the equivalent position in alphaviruses, other rubiviruses, and some bacterial macroD-type domains. In Getah virus macro this cysteine forms a covalent adduct with ADP-ribose (22). The interactions of this cysteine with ADP-ribose in the Getah virus and RuV macro structures strongly suggest that this cysteine contributes directly to catalysis in alpha- and rubiviruses, and perhaps also in some nonviral species. Another potentially important residue in the catalytic loop of RuV macro is Ser46. A serine in a similar position in Getah virus is important for ADP-ribose binding (22). A relatively small (1 Å) shift in the position of ADP-ribose relative to Ser46 would bring the ligand within hydrogen bonding distance of the Ser46 side chain. Indeed, in one of the available crystal structures of Getah virus macro, ADP-ribose is in the α-anomeric configuration and the distal ribose of ADP-ribose is rotated 60° relative to the canonical pose (22). The same rotation of the distal ribose in the RuV macro structure would reduce the distance between Ser46 and the 1’ ribose hydroxyl in the α-anomeric configuration sufficiently to form a hydrogen bond, suggesting that Ser46 and Ser30 may have analogous roles in substrate binding in RuV macro and Getah virus macro, respectively. We note that the structure of Getah virus with the unusual ribose orientation was obtained by co-crystallization with 50 mM glutamic acid. In this structure, an acetate moiety was visible in the active site positioned relative to the distal ribose so as to resemble an ADP-ribose-glutamate conjugate (22). This is consistent with the notion that MARylated glutamates are substrates for alpha- and rubivirus macro domains (15, 22, 27).

Although some macro domains can degrade PAR chains, this PAR glycohydrolase (PARG) activity requires a glutamate in the active site (14). Macros that are de-MARylases but do not degrade PAR lack a catalytic glutamate and instead rely on the acetyl group of the glutamate or aspartate protein adduct for catalysis (13). PAR hydrolysis is thus thought to proceed via a substrate-assisted mechanism, with the ADP-ribose-glutamate substrate activating the conserved structural water near the α-phosphate to nucleophilically attack the distal ribose C1’ atom (36). Consistent with its de-MARylase activity, the RuV macro has no glutamate or aspartate residues in its active site, which appears to preclude PAR degradation activity in RuV macro.

Our work demonstrates that, like alpha- and coronaviruses, rubiviruses encode a mono-ADP-ribosylhydrolase with a structurally conserved macro domain fold to counteract MARylation deposited by PARPs that are expressed upon interferon induction as part of in the host innate immune response. The macro domains from rubi-, alpha-, and coronaviruses should therefore be considered as potential targets for antiviral therapeutics or combination therapies against rubella or infections with other viruses with similar macro domains. The RuV macro crystal structures reported here could guide future efforts to develop these new therapeutics.

## MATERIALS AND METHODS

### Protein expression and purification

A gene encoding the macro domain (residues 805-983) of p150 from rubella virus strain RVi/Brooklyn.NY.USA/98/1B CRS (GenBank JN635282; UniProt G3M8F4) was cloned into the pET24a plasmid (Novagen) with a C-terminal hexahistidine purification (His_6_) tag. *Escherichia coli* BL21 (DE3) cells (New England BioLabs) were transformed with this expression construct and grown at 37°C in 2×TY medium. At an optical density (OD_600_) of 0.6-0.8, the incubator temperature was lowered to 18°C and protein expression was induced with 0.2 mM IPTG. After 16 h, the bacteria were pelleted (6,000 g for 15 min) and stored at −70°C until required. The cell pellet was resuspended in Macro-Wash Buffer (50 mM Tris pH 7.4, 0.3 M NaCl, 20 mM imidazole) supplemented with cOmplete EDTA-free protease inhibitors (Roche) and 1:10,000 (v/v) Benzonase (Sigma) and lysed by sonication. The lysate was centrifuged at 40,000 g for 30 min and the supernatant was loaded onto a HisTrap HP column (Cytiva) equilibrated in Macro-Wash Buffer. The column was washed with 30 column volumes of Macro-Wash Buffer and the protein eluted with 50 mM Tris pH 7.4, 0.3 M NaCl, 0.25 M imidazole. The protein was further purified by size-exclusion chromatography with a HiLoad 26/600 Superdex 75 pg column (Cytiva) in 20 mM HEPES pH 8, 0.15 M NaCl.

For purification of the PARP10 catalytic domain (PARP10cd), a gene encoding PARP10 residues 818-1025 (UniProt Q53GL7) codon-optimized for *E. coli* was cloned into the first multiple cloning site of pRSFDuet (Novagen) with an N-terminal glutathione S-transferase (GST) affinity tag. *Escherichia coli* BL21 (DE3) cells (New England BioLabs) transformed with this expression plasmid were grown at 37°C in 2×TY medium, induced with 0.2 mM IPTG at OD_600_ = 0.6-0.8, and incubated at 18°C for 16-18h. The cells were pelleted and resuspended in PARP-Wash Buffer (50 mM Tris pH 8.0, 0.2 M NaCl, 0.1 mM EDTA, 10% glycerol, 1 mM DTT) supplemented with cOmplete EDTA-free protease inhibitors (Roche) and 1:10,000 (v/v) Benzonase (Sigma). Cells were lysed by sonication. The lysate was centrifuged at 40,000 g for 30 min and the supernatant loaded onto a GST-Trap 4B column (Cytiva) equilibrated in PARP-Wash Buffer. The column was washed with 30 column volumes of PARP-Wash Buffer and the protein eluted with PARP-Wash Buffer containing 25 mM reduced glutathione (GSH). The protein was further purified by size-exclusion chromatography on a Superdex S200 10/300 Increase column (Cytiva) in 20 mM HEPES pH 8.0, 0.2 M NaCl, 10% glycerol, and 0.5 mM tris(2-carboxyethyl)phosphine hydrochloride (TCEP).

### Crystallographic structure determination

Crystals were grown at 18°C by sitting drop vapor diffusion. The purified protein was concentrated to 9 g L^-1^ (452 µM) and mixed with an equal volume of reservoir solution optimized from the PEG/LiCl Grid Screen (Hampton Research): 14.4% PEG 6K, 1.26 M LiCl, 0.1 M Bicine pH 9. Crystals were cryoprotected in reservoir solution supplemented with 10% glycerol, and frozen in liquid nitrogen. For the ADP-ribose-bound structure, crystals were soaked in mother liquor supplemented with 2 mM ADP-ribose for 30-45 min prior to freezing. X-ray diffraction data were collected at 100 K at Diamond Light Source beamlines I03 and I04. Data were processed with autoPROC v1.0.5 (41), and STARANISO v1.0.4 (42). Molecular replacement was performed with MrBUMP (34) as implemented in CCP4 (35) using an atomic search model based on the human protein-proximal ADP-ribosyl-hydrolase MacroD2 structure (PDB 4IQY) (36). An initial model was built using AutoBuild in PHENIX v1.20.1 (43), manually completed with COOT (44) and iteratively refined with PHENIX. See Table 1 for crystallographic data collection, refinement, and validation statistics.

### Isothermal titration calorimetry (ITC)

Binding of RuV macro to ADP-ribose was analyzed in 20 mM HEPES pH 8, 0.15 M NaCl at 25°C, with an AutoiTC200 calorimeter (MicroCal). The sample cell was loaded with 0.2 ml of 100 µM RuV macro and the titrant syringe with 2 mM ADP-ribose. 20 serial injections of 2 μl ADP-ribose were performed at 3 min intervals. The stirring speed was 1,000 rpm and the reference power was maintained at 6.06 μcal/s. The net heat absorption or release associated with each injection was calculated by subtracting the heat associated with the injection of ADP-ribose to buffer. Thermodynamic parameters were extracted from a curve fit to the data to a single-site model with Origin 7.0 (MicroCal). Experiments were performed in duplicate.

### Differential scanning fluorimetry (DSF)

10 µl samples of 25 µM RuV macro in 20 mM HEPES pH 8, 0.15 M NaCl, with or without addition of 5 mM ADP-ribose, were loaded into glass capillaries (NanoTemper) by capillary action. Intrinsic protein fluorescence at 330 nm and 350 nm, F330 and F350, respectively, was measured from 15°C to 95°C with a ramp rate of 2°C per minute with a Prometheus NT.48 nano-fluorimeter (NanoTemper). The melting temperatures (T_m_ values) were calculated with the accompanying software (NanoTemper) as the temperature at the peak of the first derivative of F350:F330 versus temperature.

### Enzymatic de-ADP-ribosylation assay

Following a previously established protocol (23), auto-mono-ADP-ribosylated PARP10cd was obtained by incubating 10 μM purified PARP10cd in 1 mM NAD+ for 20 min at 37°C in Reaction Buffer (50 mM HEPES pH 8.0, 0.15 M NaCl, 0.2 mM TCEP, 0.02% NP-40). The de-ADP-ribosylation reaction was performed by incubating purified RuV macro with mono-ADP-ribosylated PARP10cd at 37°C in Reaction Buffer at equimolar ratios (1 μM). The reaction was terminated at 0, 1, 2, 4, 8, 16, 32 or 64 min by adding SDS-PAGE sample loading buffer and incubating at 95° for 5 min. Following separation SDS-PAGE on a 4-20% gradient gel, the proteins were transferred to a PVDF membrane and the de-MARylation activity determined by Western blotting. The membrane was blocked in 5% milk in PBS, 0.05% Tween-20. The membrane was incubated for 3 h at 22°C in Anti-mono-ADP-Ribose Binding Reagent (Sigma, MABE1076, RRID:AB_2665469; 1:3,000 dilution) in place of a primary antibody. The secondary antibody was anti-rabbit immunoglobin (Cell Signaling Technology, 5151S, RRID:AB_10697505, DyLight® 800; dilution 1:10,000 dilution, 30 min at 22°C). Blots were imaged with the near-infrared system of an Odyssey fluorescent scanner (LI-COR Biosciences) after washing with PBS and water.

### Data availability

The atomic coordinates of the rubella p150 macro domain with and without ADP-ribose were deposited in the Protein Data Bank with accession codes 8P0C [https://doi.org/10.2210/pdb8p0c/pdb] and 8P0E [https://doi.org/10.2210/pdb8p0e/pdb].

## Acknowledgements.

Crystallographic data were collected on beamlines I03 and I04 at Diamond Light Source (DLS). Access to DLS (proposal MX21426) was supported by the Wellcome Trust, MRC and BBSRC. We thank Fabrice Gorrec (MRC-LMB Crystallization Facility) for advice on protein crystallization. We thank Dom Bellini (MRC-LMB X-ray Crystallography Facility) for advice in crystallographic data collection. This work was supported by MRC research grant MR/S021604/1 to Y.M., Wellcome Trust Senior Research Fellowship 217191/Z/19/Z to Y.M., and Wellcome Trust PhD Studentship 205833/Z/16/Z to G.S. Open access publication was funded by the University of Cambridge.

